# Extensive epitranscriptomic methylation of A and C residues on murine leukemia virus transcripts enhances viral gene expression

**DOI:** 10.1101/591545

**Authors:** David G. Courtney, Andrea Chalem, Hal P. Bogerd, Brittany A. Law, Edward M. Kennedy, Christopher L. Holley, Bryan R. Cullen

**Author notes:** Oncorus, 50 Hampshire Street, Suite 401, Cambridge MA 02139.

## Abstract

While it has been known for several years that viral RNAs are subject to the addition of several distinct covalent modifications to individual nucleotides, collectively referred to as epitranscriptomic modifications, the effect of these editing events on viral gene expression has been controversial. Here, we report the purification of murine leukemia virus (MLV) genomic RNA to homogeneity and show that this viral RNA contains levels of *N*^6^-methyladenosine (m^6^A), 5-methylcytosine (m^5^C) and 2’*O*-methylated (Nm) ribonucleotides that are an order of magnitude higher than detected on bulk cellular mRNAs. Mapping of m^6^A and m^5^C residues on MLV transcripts identified multiple discrete editing sites and allowed the construction of MLV variants bearing silent mutations that removed a subset of these sites. Analysis of the replication potential of these mutants revealed a modest but significant attenuation in viral replication in 3T3 cells in culture. Consistent with a positive role for m^6^A and m^5^C in viral replication, we also demonstrate that overexpression of the key m^6^A reader protein YTHDF2 enhances MLV replication, while downregulation of the m^5^C writer NSUN2 inhibits MLV replication.

**Importance:** The data presented in this manuscript demonstrate that MLV RNAs bear an exceptionally high level of the epitranscriptomic modifications m^6^A, m^5^C and Nm, thus suggesting that these each facilitate some aspect of the viral replication cycle. Consistent with this hypothesis, we demonstrate that mutational removal of a subset of these m^6^A or m^5^C modifications from MLV transcripts inhibits MLV replication in *cis* and a similar result was also observed upon manipulation of the level of expression of key cellular epitranscriptomic cofactors in *trans*. Together, these results argue that the addition of several different epitranscriptomic modifications to viral transcripts stimulates viral gene expression and suggest that MLV has therefore evolved to maximize the level of these modifications that are added to viral RNAs.

## Introduction

Eukaryotic mRNAs are subject to a range of covalent modifications at the single nucleotide level and it is now evident that these epitranscriptomic modifications can profoundly affect mRNA function (1–3). While the most prevalent epitranscriptomic mRNA modification involves methylation of the *N*^6^ position of adenosine (m^6^A), several other mRNA modifications, including cytidine methylation (m^5^C) and 2’*O*-methylation of the ribose moiety that forms part of all four ribonucleotides (Am, Gm, Cm and Um, collectively Nm), have also been reported.

Addition of m^6^A is the most intensively studied epitranscriptomic modification and the protein complex responsible for m^6^A addition, or m^6^A “writer”, has been identified as a nuclear heterotrimer, consisting of the proteins METTL3, METTL14 and WTAP, that adds m^6^A to mRNA sites bearing the consensus sequence 5’-RA*C-3’, where R is a purine (1–3). Once added, m^6^A can be recognized by several “readers”, including the nuclear YTHDC1 and cytoplasmic YTHDF2 proteins, which then regulate the splicing, translation and/or stability of that mRNA. Less is known about the m^5^C modification, although NSUN2 has been shown to add m^5^C to a handful of cellular mRNAs (4–7) and we have recently identified NSUN2 as the primary writer of m^5^C residues on the HIV-1 genome (8).

Previously, we reported that m^6^A residues enhance viral gene expression and replication for HIV-1, influenza A virus and the polyoma virus SV40 (9–11) and others have also reported that m^6^A residues promote HIV-1 and enterovirus 71 replication (12, 13) and play a role in the activation of lytic replication in Kaposi’s sarcoma herpesvirus (KSHV)-infected cells (14, 15). More recently, we reported that addition of m^5^C also enhances HIV-1 gene expression (8), and others have reported that Nm modifications on HIV-1 transcripts promote HIV-1 replication by inhibiting the detection of viral transcripts by the innate antiviral RNA sensor MDA5 (16). Consistent with a positive role for these epitranscriptomic modifications in the regulation of viral replication, we recently reported that HIV-1 transcripts bear a far higher level of m^6^A, m^5^C and Nm residues than does the average cellular mRNA (8). Here, we extend these earlier findings by demonstrating that the addition of both m^6^A and m^5^C independently upregulates murine leukemia virus (MLV) gene expression and replication. Moreover, we further show that MLV transcripts are also extensively epitrancriptomically modified, with m^6^A, m^5^C and Nm residues all detected at levels that range from 7 to >20-fold higher than observed on cellular poly(A)+ RNA. Together, these observations are consistent with the hypothesis that sites of epitranscriptomic modification on viral mRNAs are under positive selection and suggest that many viruses may have evolved to use epitranscriptomic gene regulation as a mechanism to promote their replication and, hence, pathogenesis.

## Results

### Extensive epitranscriptomic modification of the MLV RNA genome

The initial goal of this project was to quantify the epitranscriptomic RNA modifications present on MLV genomic RNA (gRNA) using ultra-high-performance liquid chromatography and tandem mass spectrometry (UPLC-MS/MS) (17) (8). This required the purification of the gRNA away from cellular tRNAs and other ncRNAs that are heavily modified. MLV virions released into the supernatant media from 3T3 cells infected with MLV derived from the pNCS proviral plasmid (18) were pelleted through a 20% sucrose cushion followed by banding on a 7.2% to 20% iodixanol gradient, which separates retroviral virions from cellular exosomes and debris (19). Because MLV virions contain high levels of tRNAs, 7SL RNA and other cellular ncRNAs (20), isolation of MLV virion particles is necessary but not sufficient to yield pure MLV gRNA. Therefore, we next isolated total virion RNA, denatured it in urea loading dye and then size fractionated the RNA by electrophoresis on a 1.5% TBE preparative agarose gel. The ~8kb MLV gRNA was then visualized and excised. This procedure was performed in triplicate to yield three independent MLV gRNA preparations.

To assess the purity of the MLV gRNA preparations, we performed RNA-seq and then aligned the reads obtained first to the mouse genome and then to the MLV genome. As shown in Fig. 1A, 99.48% of the reads obtained from gRNA preparation 1 aligned to the MLV genome, while only 0.52% aligned to the mouse genome, and closely similar results were obtained for MLV gRNA preparations 2 and 3. These data also revealed that the MLV-specific reads obtained aligned to the entire MLV genome (Fig. 1B). We next quantified the precise level of several epitranscriptomic modifications on the MLV RNA genome using UPLC-MS/MS, as previously described (17). Quantification of the level of multiple epitranscriptomic modifications across the three MLV gRNA preparations revealed a high level of reproducibility (Figs. 1C and 1D). These data also revealed a particularly high level of m^6^A (~51 m^6^A residues per gRNA) as well as high levels of 2’*O*-methylated adenosine (Am), guanosine (Gm) and cytosine (Cm) (~27, ~25 and ~18 residues, respectively, per gRNA), as well as a high level of m^5^C (~13 residues per gRNA). These levels are considerably higher than previously reported for cellular poly(A)+ RNA (2, 21) (8). Specifically, the level of m^6^A reported for both human and murine poly(A)+ RNA is ~0.35% of “A” residues, while ~2.4% of all “A” residues in the MLV genome are m^6^A (Fig. 1D). It has also been reported that ~0.06% of all “C” residues in poly(A)+ RNA are modified to m^5^C, compared to ~0.54% of “C” residues in the MLV gRNA (Fig. 1D). Similarly, Am, Gm and Cm represent from 0.74% to 1.25% of their cognate residue in the MLV genome, while we have previously reported that these 2’*O*-Me-modified nucleotides each represent <0.1% of the A, G and C residues present on cellular poly(A)+ RNA (8). We therefore conclude that m^6^A, m^5^C, Am, Gm and Cm are all highly overrepresented on MLV gRNAs, when compared to the average cellular mRNA.

**Figure 1.**
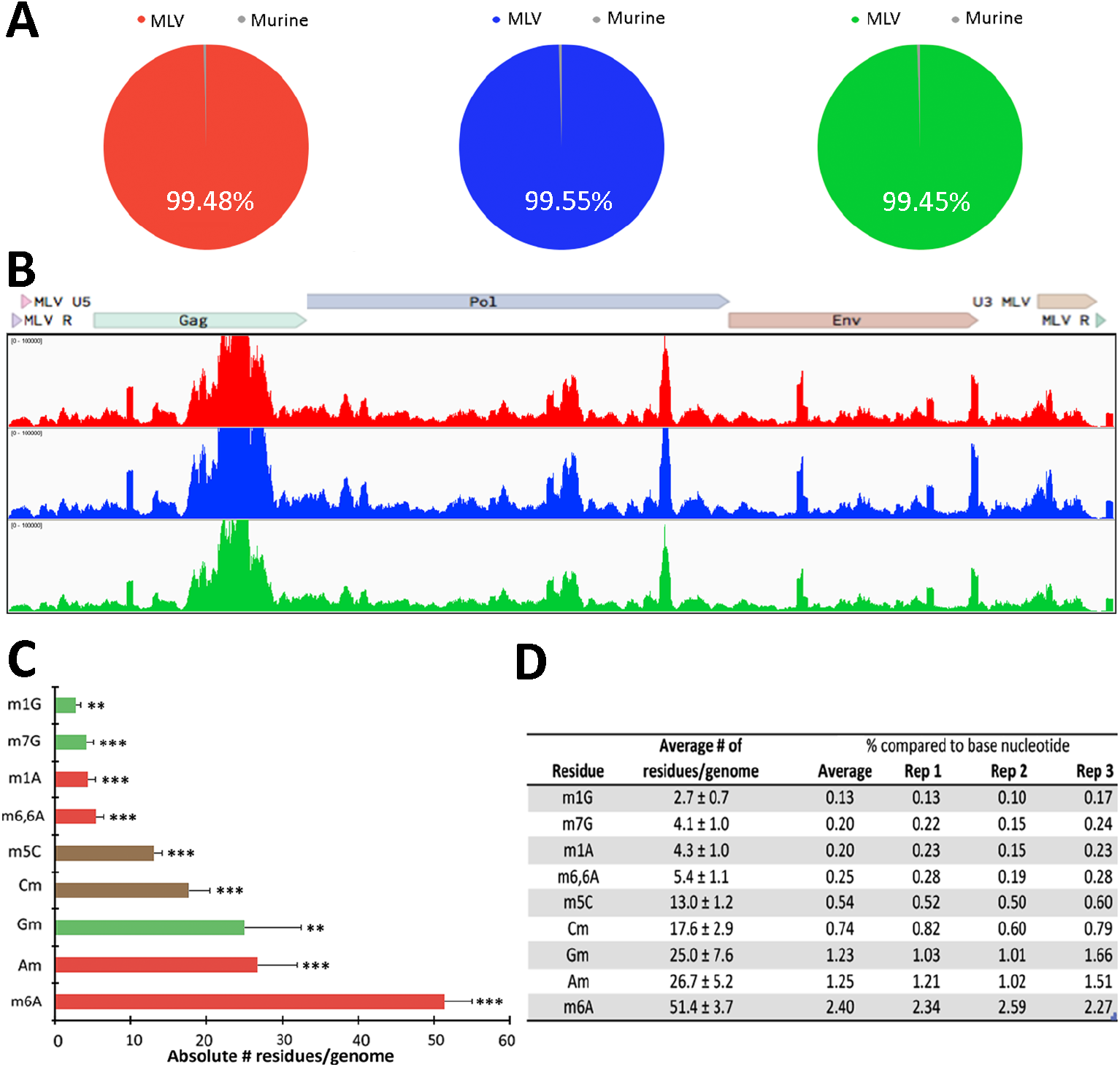
Extensive epitranscriptomic modifications of MLV gRNAs. (A) Alignment of the RNA-seq reads obtained from the three MLV gRNA preparations to the MLV or mouse genome. (B) Alignment of RNA-seq reads from the three MLV gRNA preparations to the MLV genome, demonstrating coverage of the entire MLV genome. (C) Quantification of the absolute number of nine different RNA modifications on MLV gRNA, as determined by UPLC-MS/MS analysis of three independent gRNA samples, with SD indicated. **, p<0.01; ***, p<0.001. (D) Table of the UPLC-MS/MS data described in panel C, showing the percentage abundance of each modification in comparison to the parental nucleotide for each replicate, with good concordance between samples.

While our data identify the five epitranscriptomic modifications listed above as unusually prevalent on MLV gRNA, with >10 modified residues of each per gRNA, we also detected several other modified nucleotides at levels ranging from ~2.7 to ~5.4 residues per MLV genome (Figs. 1C and 1D). In the case of 1-methylguanosine (m^1^G) and 7-methylguanosine (m^7^G), the observed levels in the MLV genome are comparable to levels detected previously in cellular poly(A)+ RNA (22) (8). However, this is not true for *N^6^,N^6^*-dimethyladenosine (m^6,6^A), which represents ~0.25% of all A residues in the MLV gRNA, versus ~0.037% in total poly(A)+ RNA, a difference of ~7-fold. Similarly, 1-methyladenosine (m^1^A) was detected at an ~22-fold higher level on MLV gRNAs than detected on cellular poly(A)+ RNA (0.20% vs. 0.009%) (8). However, as these four residues are all present at very low levels on MLV gRNAs (Fig. 1C), it is unclear whether they exert any phenotypic effect. While we did not detect any epitranscriptomically modified “U” residues, such as 2’*O*-Me-uridine (Um) or pseudouridine, we note that the UPLC-MS/MS method is less sensitive for detecting modified uridines at low concentrations of input RNA, as in this case, due to the comparatively inefficient ionization of uridine compared to other ribonucleosides. Nevertheless, our data do suggest that levels of Um in the MLV gRNA are <1nM, which equates to <20 Um residues per MLV gRNA.

### Mapping of m^6^A and m^5^C residues on the MLV RNA genome

Having demonstrated that MLV gRNAs bear a substantial number of m^6^A and m^5^C residues, we wished to map these residues not only on the gRNA but also on MLV RNAs expressed in infected 3T3 cells. For this purpose, we infected 3T3 cells with MLV virions rescued from the pNCS proviral clone (18) and, at 48 hours post-infection (hpi), pulsed the cells with the highly photoreactive uridine analog 4-thiouridine (4SU) for a further 24 h (23). We then isolated MLV virions, as described above, from the supernatant media of MLV-infected 3T3 cells and purified total virion RNA. In parallel, we also harvested total RNA from MLV-infected 3T3 cells and subjected this RNA to a single round of poly(A)+ isolation to enrich for mRNAs. The MLV virion and MLV-infected cell RNA preparations were then subjected to the previously described PA-m^6^A-seq or PA-m^5^C-seq procedures (24) (8). Briefly, the purified 4SU-labeled RNAs were incubated with an antibody specific for either m^6^A or m^5^C and then UV-irradiated to crosslink the antibody to the bound site. The resultant RNA:protein complexes were then incubated with RNase T1, to remove unbound RNA, and the bound ~20 nt RNA fragments recovered, converted to cDNA and subjected to deep sequencing. As may be observed in Fig. 2A, we detected several m^6^A peaks on the MLV gRNA, almost all of which were also observed on the MLV transcripts expressed in infected 3T3 cells. Consistent with the fact that all MLV virion RNAs are genome length, while ~50% of the MLV transcripts expressed in infected cells are *env* mRNAs that have been spliced to remove the Gag and Pol open reading frames (ORFs), we detected a 2-3-fold lower level of m^6^A binding sites in the MLV *gag* and *pol* genes, when compared to the *env* gene and LTR, in the intracellular RNA sample (Fig. 2A). While our UPLC-MS/MS data indicate that each MLV gRNA contains ~51 m^6^A residues (Fig. 1D), we only detected ~20 major m^6^A sites using the PA-m^6^A-seq technique. While the reasons for this discrepancy are unclear, we note that it has been recently reported that m^6^A residues embedded in duplex RNA are not readily detected by m^6^A-specific antibodies (25). Nevertheless, this discrepancy does suggest that the m^6^A sites that were detected on the MLV RNA genome are likely to be heavily modified.

**Figure 2.**
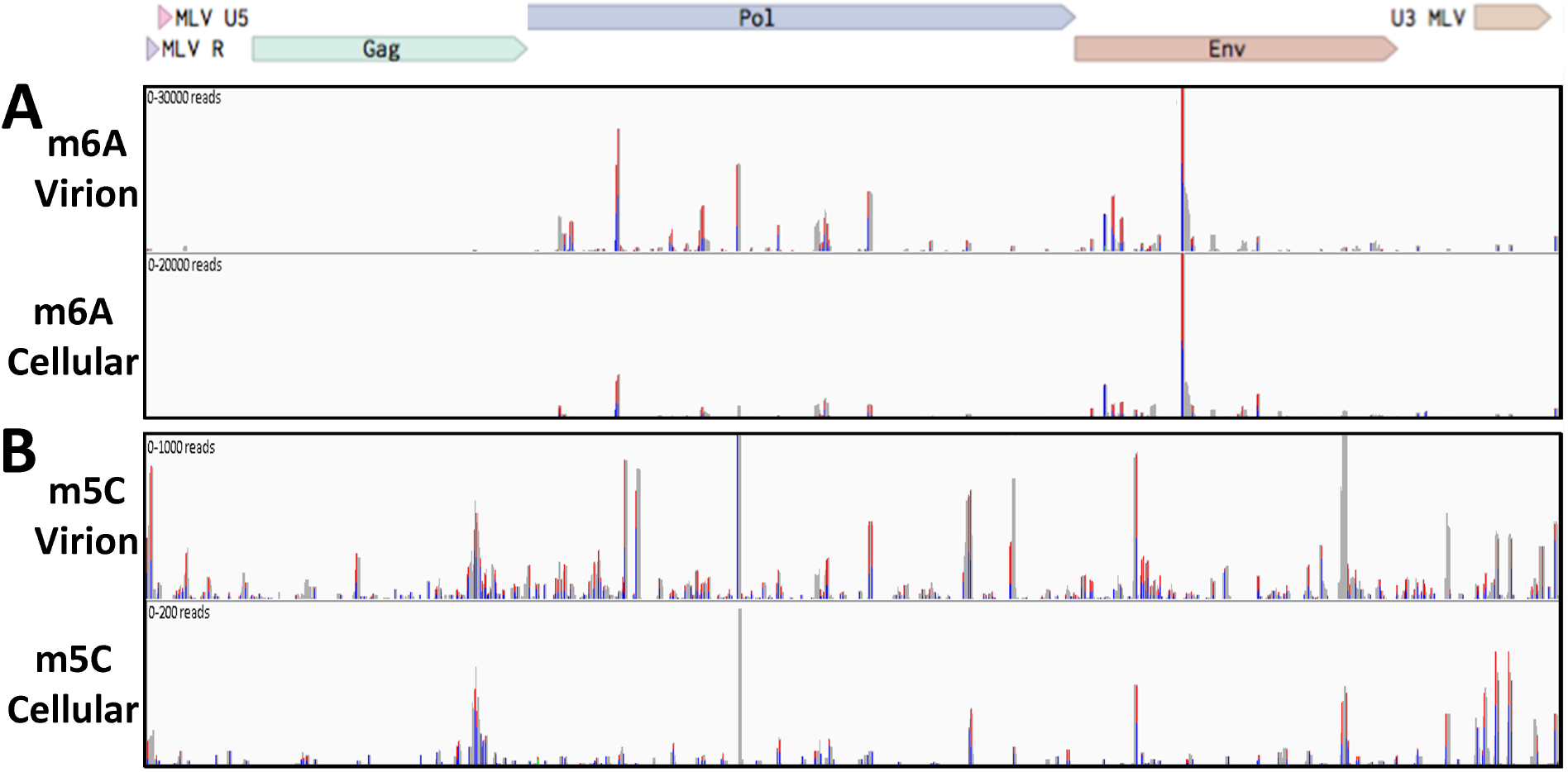
Mapping of m^6^A and m^5^C residues on infected cell and virion-derived MLV RNAs. (A) The m^6^A residues located on MLV gRNA isolated from virions (top lane), or from MLV RNAs isolated from infected 3T3 cells (bottom lane), were mapped using the antibody-based PA-m^6^A-seq technique. (B) The m^5^C residues on MLV gRNA isolated from virions (top lane), or from viral transcripts expressed in MLV-infected 3T3 cells (bottom lane), were mapped using the antibody-based PA-m^5^C-seq technique. Blue peaks, single T to C conversion, red peaks, more than one T to C conversion.

The PA-m^5^C-seq technique also mapped a substantial number of m^5^C residues on the MLV genome and, as expected, revealed minimal overlap with the mapped m^6^A sites (Fig. 2B). We again detected a higher level of antibody binding in the gag/pol region of the MLV genome in the virion-derived RNA sample than in the intracellular MLV RNA, although this varied somewhat by peak. In contrast to the PA-m^6^A-seq data, which identified somewhat fewer m^6^A modification sites on the MLV gRNA than predicted by the UPLC-MS/MS data, the PA-m^5^C-seq data detected ~40 m^5^C sites on the MLV gRNA (Fig. 2B), which is more than the ~13 sites predicted by the UPLC-MS/MS technique (Fig. 1D), thus suggesting that most of these m^5^C sites are likely to be only partially modified.

### Epitranscriptomic addition of m^6^A and m^5^C facilitates MLV gene expression

One way to test whether the addition of m^6^A or m^5^C has any effect on MLV gene expression and replication is to mutate the locations of these modifications by, for example, changing mapped m^6^A residues to “G” residues and mapped m^5^C residues to “U” residues. In the case of m^6^A, the modified “A” is found in the context of the sequence 5’-RA*C-3’, where R is a purine (1), so modified “A” residues are easier to identify within the ~20 nt peaks mapped in Fig. 2A. Nevertheless, many peaks do contain more than one 5’-RA*C-3’ consensus sequence. In contrast, m^5^C modifications on mRNAs do not occur in a sequence consensus (8), so there are generally multiple “C” residues within each of the m^5^C peaks mapped in Fig. 2B. This complicates the design of MLV mutants lacking specific m^6^A or m^5^C sites, as interpretation of the results obtained requires that all the introduced mutations are silent, which in practice means located in the wobble position of codons.

In the case of m^6^A, it proved impossible to design silent mutations that would ablate most of the m^6^A sites mapped in Fig. 2A, though this was possible for all three of the 5’-RA*C-3’ motifs found in one of the most prominent peaks, in the MLV *env* gene, to generate the MLV-Δm^6^A mutant (indicated by # in Fig. 3A, lane 1). In the case of m^5^C, we were fortunate that four prominent m^5^C peaks located in the MLV *pol* gene (indicated by # in Fig. 3A, lane 3) could be silently mutated in the context of the pNLS proviral expression vector to generate the MLV-Δm^5^C mutant.

**Figure 3.**
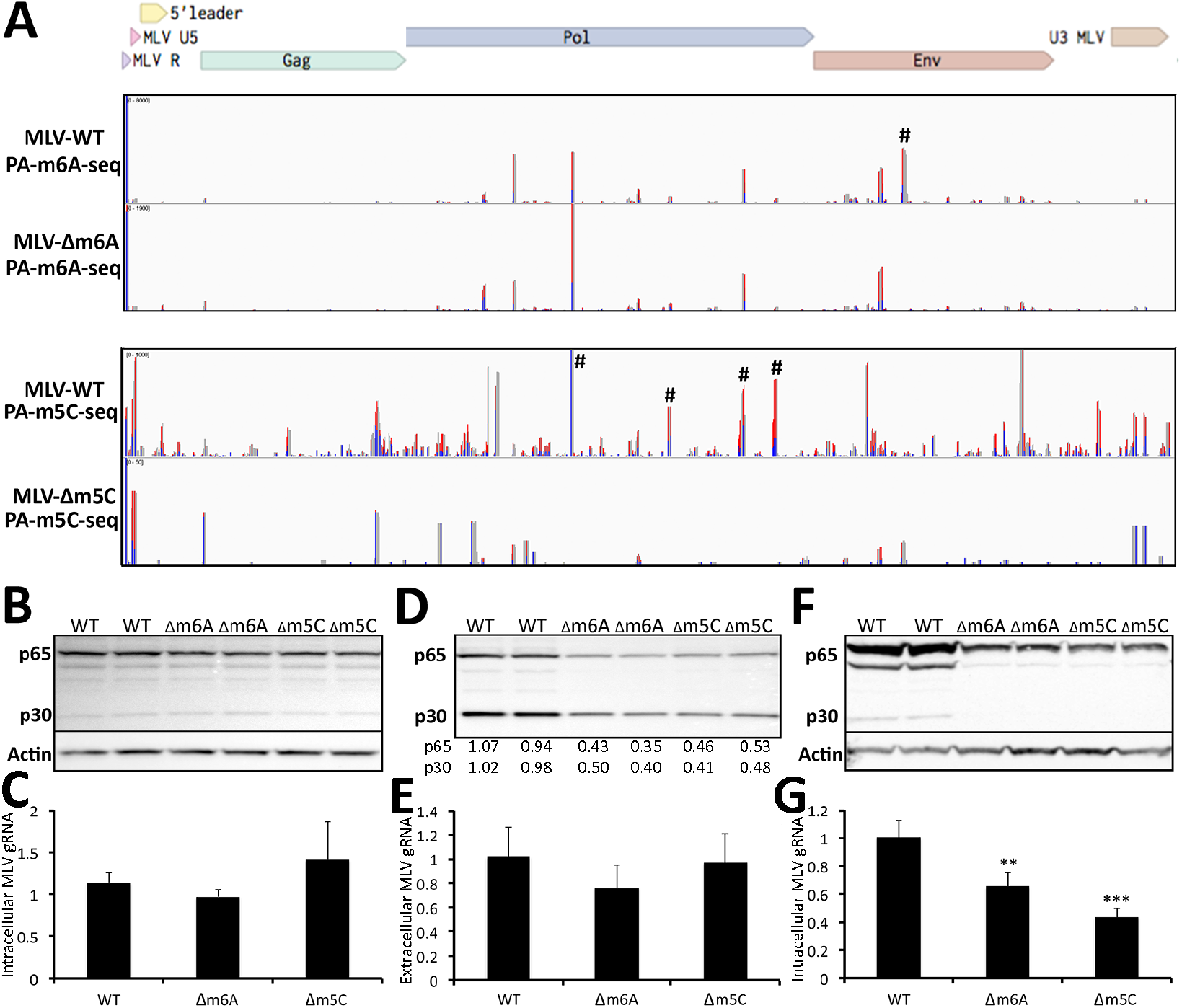
Loss of m^6^A or m^5^C residues on MLV RNAs reduces MLV gene expression. (A) Alignment of PA-m^6^A-seq reads to intracellular MLV RNA isolated from 3T3 cells infected with wild-type MLV (first lane) or the MLV-Δm^6^A mutant (second lane), Similarly, this figure also shows an alignment of PA-m^5^C-seq reads to intracellular MLV RNAs expressed in 3T3 cells infected with wild-type MLV (third lane) or MLV-Δm^5^C (fourth lane). # denotes peaks where silent mutations were introduced to ablate specific m^6^A or m^5^C addition sites. (B) Western blot of the MLV Gag proteins p65 and p30 expressed from wild type MLV, MLV-Δm^6^A or MLV-Δm^5^C in 293T cells transfected with wildtype or mutant pNCS-based plasmids, at 72 h post-transfection. Representative assays are shown in duplicate. (C) qPCR of MLV RNA in total RNA isolated from 293T cells transfected with wild type MLV, MLV-Δm^6^A or MLV-Δm^5^C at 72 h post-transfection, normalized to GAPDH mRNA. Average of three independent experiments with SD indicated. (D) Western blot of the MLV Gag proteins p65 and p30 from virions isolated from equal amounts of the supernatant media from 293T cells expressing wild type MLV, MLV-Δm^6^A or MLV-Δm^5^C, as shown in panel B. Band intensities were quantified by ImageJ and normalized to the average level seen with wild type MLV, with numbers given below the panel. Representative assays are shown in duplicate. (E) qPCR quantification of MLV gRNA prepared from virions isolated from the supernatant media of 293T cells expressing wild type MLV, MLV-Δm^6^A or MLV-Δm^5^C, and normalized to 7SL RNA. These are the same virions analyzed in panel D. Average of three independent experiments with SD indicated. (F) Western blot of the MLV Gag proteins p65 and p30 isolated from 3T3 cells infected with equal amounts of wild type MLV, MLV-Δm^6^A or MLV-Δm^5^C, as determined in panel D, at 72 hpi. Representative assays are shown in duplicate. (G) qPCR quantification of MLV gRNA from the same 3T3 cells shown in panel F, infected with equal amounts of wild type MLV, MLV-Δm^6^A or MLV-Δm^5^C MLV, normalized to cellular GAPDH mRNA. Average of three independent experiments with SD indicated. **, p<0.01; ***, p<0.001.

To confirm that these introduced mutations indeed resulted in the loss of the predicted modified residues, we rescued the pNCS-based MLV-Δm^6^A and MLV-Δm^5^C mutants by transfection into 293T cells followed by culture in susceptible 3T3 cells. We then performed PA m^6^A-seq and PA-m^5^C-seq using intracellular RNA preparations derived from 4SU-pulsed 3T3 cells infected with wild type MLV, or the MLV-Δm^6^A or MLV-Δm5C mutants. As shown in Fig. 3A (upper two panels), the mutations introduced into the MLV-Δm^6^A mutant, indicated by #, resulted in the precise loss of the predicted major m^6^A peak, while other peaks were unaffected. In the case of the MLV-Δm^5^C mutant, the four introduced mutations not only ablated the four targeted peaks in the MLV *pol* gene (indicated by # in the lower two panels of Fig. 3A) but also seemed to result in an overall reduction in m^5^C addition, even at sites that were not altered. The reasons for this effect are not clear, but it could imply that m^5^C addition to RNAs, unlike m^6^A addition, is in some way cooperative.

Next, we assessed whether the introduced mutations affected MLV gene expression and/or replication. For this purpose, we transfected 293T cells with the wild type MLV proviral expression vector pNCS, or with the pNCS-Δm^6^A or pNCS-Δm^5^C mutant plasmids. We detected comparable levels of MLV Gag production in the transfected 293T cells (Fig. 3B), as well as similar levels of MLV gRNA expression (Fig. 3C). However, analysis of the supernatant media revealed that the MLV-Δm^6^A mutant released 2-3-fold less MLV Gag protein into the supernatant media, and 293T cells transfected with the MLV-Δm^5^C mutant also released ~2-fold less MLV Gag protein (Fig. 3D). One possibility we considered is that the mutations present in the MLV-Δm^6^A or MLV-Δ m^5^C mutant might affect gRNA packaging into MLV virion particles. To address this, we performed qRT-PCR to measure the level of MLV gRNA isolated from the supernatant media of the transfected 293T cells and then normalized these data by qRT-PCR analysis of the level of cellular 7SL RNA, which is known to also be selectively packaged into MLV virions (20). As shown in Fig. 3E, this analysis did not reveal any reduced packaging into virions of the MLV gRNA produced by the MLV-Δm^6^A or MLV-Δm^5^C mutants, although clearly fewer virions were released (Fig. 3D).

Next, we normalized the MLV-containing supernatants obtained from the transfected 293T cells, using the MLV Gag quantitations shown in Fig. 3D, and then used equal amounts of each MLV variant to infect susceptible 3T3 cells. At 72 hpi, we harvested these infected cells and analyzed MLV Gag protein expression (Fig. 3F) and gRNA expression (Fig. 3G). As may be observed, we detected a significant reduction in the level of both Gag protein and MLV gRNA in the cultures infected with the MLV-Δm^6^A and MLV-Δm^5^C mutant, though this effect was only 2-3-fold. We note, however, that as these mutants retain a substantial number of m^6^A and m^5^C sites, a modest phenotype is not unexpected. Nevertheless, these data are clearly consistent with the hypothesis that the m^6^A and m^5^C epitranscriptomic modifications detected on MLV transcripts are both able to facilitate some aspect of MLV replication.

While the mutations introduced into the MLV-Δm^6^A and MLV-Δm^5^C mutants are designed to be silent and to not impact sequences with a known regulatory role, it remains possible that they could affect an important RNA structure or protein binding site unrelated to the targeted RNA modifications. We therefore wished to test whether inhibition of m^6^A or m^5^C addition, or their enhanced detection by cellular readers, might also impact MLV gene expression. Previously, we reported that overexpression of the key cellular m^6^A reader YTHDF2 increases viral gene expression for three distinct viral species, viz. HIV-1, influenza A virus and the SV40 (9–11), and we therefore asked whether stable overexpression of murine YTHDF2 in 3T3 cells would also enhance MLV gene expression. For this purpose, we generated clonal 3T3 cell lines transduced with a lentiviral vector expressing either FLAG-tagged YTHDF2 or GFP and selected single cell clones expressing readily detectable levels of these proteins (Fig. 4A). We then infected these cells with wild type MLV and assessed viral gene expression at 48 and 72 hpi by Western blot for MLV p65 Gag. As shown in Fig. 4A, YTHDF2 overexpression indeed resulted in a readily detectable increase in MLV Gag expression, thus again arguing that m^6^A addition to MLV transcripts facilitates viral gene expression and replication.

**Figure 4.**
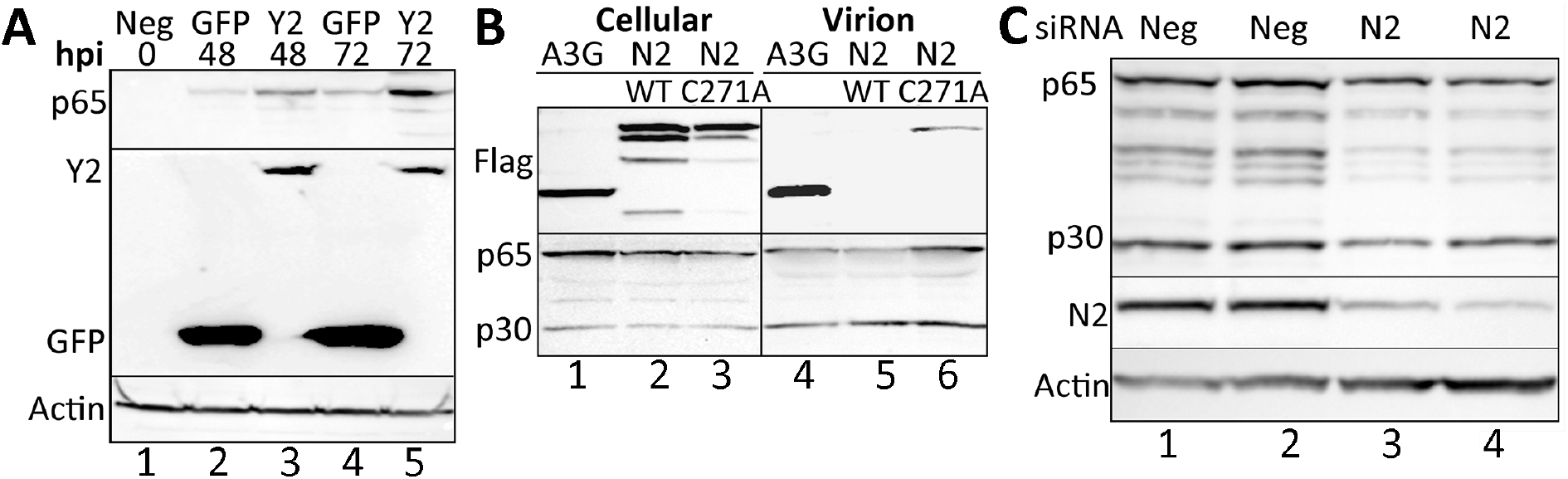
Alteration of m6A or m5C machinery affects MLV protein levels. (A) Stable overexpression of murine YTHDF2 in 3T3 cells increases MLV Gag protein expression at both 48 and 72 hpi, when compared to control, GFP-overexpressing 3T3 cells. Y2; YTHDF2. (B) Transient overexpression of APOBEC3G (A3G), wild type NSUN2 (N2) or NSUN2-C271A in MLV-expressing 293T cells. All three overexpressed proteins are present in the intracellular lysate but only A3G and the mutant NSUN2-C271A are detectably packaged into MLV virions. A representative experiment is shown. (C) siRNA knockdown of NSUN2 (N2) in 293T cells expressing full-length MLV reduces the expression of the MLV Gag proteins. Representative assays are shown in duplicate.

While the proteins that “write” and “read” m^6^A modifications are well defined, this is less clear for the m^5^C modification as several cytidine methyltransferases have been described. However, the cellular protein NSUN2 has previously been reported to add m^5^C to specific cellular mRNAs (4–7) and we have recently reported that NSUN2 is primarily responsible for the addition of m^5^C modifications to HIV-1 transcripts (8). An interesting aspect of NSUN2 is that it forms a transient covalent bond with the “C” residues it is methylating and release requires the action of a conserved cysteine residue located at position 271 in NSUN2. As a result, mutagenesis of cysteine 271 to alanine (C271A) leads to the spontaneous formation of NSUN2 crosslinks to target “C” residues on RNAs (6). Therefore, in cells expressing NSUN2-C271A, we predicted that this mutant protein would crosslink to MLV gRNA and potentially be packaged into MLV virions. As shown in Fig. 4B, we indeed observed packaging of the NSUN2-C271A mutant, but not wild type NSUN2, into MLV virions produced in transfected 293T cells, thus not only identifying NSUN2 as an enzyme that adds m^5^C to MLV gRNAs but also, more generally, confirming that MLV gRNAs do indeed bear “C” residues that are methylated in producer cells.

We next asked in reduced expression of NSUN2 would result in a reduction in MLV gene expression. This was achieved by the efficient knockdown of NSUN2 expression using RNA interference (RNAi), as shown in Fig. 4C. Importantly, knockdown of NSUN2 expression in 3T3 cells also resulted in a marked drop in the expression of the MLV Gag proteins (Fig. 4C). Therefore, consistent with the data presented in Fig. 3, these results argue that addition of m^5^C, like addition of m^6^A, to MLV transcripts enhances MLV RNA expression.

## Discussion

Previously, we and others have reported that the addition of m^6^A facilitates viral gene expression and replication for a range of distinct viruses, including HIV-1, influenza A virus, SV40, enterovirus 71 and KSHV (9–15). More recently, we have also presented evidence indicating that m^5^C promotes HIV-1 mRNA translation and gene expression (8), while others have reported that Nm residues on HIV-1 transcripts promote HIV-1 replication by inhibiting activation of the innate antiviral RNA sensor MDA5 (16). Together, these observations indicate that at least a subset of the epitranscriptomic modifications found on mRNAs, specifically m^6^A, m^5^C and Nm, each acts as a positive regulator of some aspect of the viral replication cycle and should therefore be selected for during viral evolution. Consistent with this hypothesis, we recently reported that m^6^A, m^5^C and Nm ribonucleotides were all present at substantially higher levels on the HIV-1 RNA genome than on poly(A)+ RNA isolated from human cells, with m^5^C (~20x higher) and Nm (11-28x higher) being particularly enriched (8).

Despite the evidence delineated above arguing for a positive role for at least some epitranscriptomic modifications in the viral life cycle, this issue has remained controversial. Specifically, one group has argued that m^6^A actually inhibits HIV-1 gene replication (26) and others have suggested that m^6^A modification of flaviviral transcripts, including Zika virus RNAs, inhibited some aspect of the viral replication cycle (27, 28). Why a rapidly evolving, lytically replicating virus, such as Zika virus, should retain m^6^A sites if these inhibit virus replication in *cis* was not, however, investigated.

In this manuscript, we have sought to shed further light on the role of epitranscriptomic modifications in regulating viral gene expression and replication, and we present three lines of evidence arguing that m^6^A, m^5^C and Nm ribonucleotides indeed exert a positive effect on the replication of the retrovirus MLV when present in *cis* on viral RNAs. Firstly, we demonstrate that MLV genomic RNAs are subject to exceptionally high levels of modification by addition of m^6^A, m^5^C and Nm, with observed levels that are from 7 to >20-fold higher than observed in cellular poly(A)+ RNA (Fig. 1). Secondly, we mapped several sites of m^6^A and m^5^C addition on MLV transcripts and then mutationally ablated a small number of these by the introduction of silent mutations (Figs. 2 and 3). While these two mutant viruses, termed MLV-Δm^6^A and MLV-Δm^5^C, retained the majority of their m^6^A and m^5^C residues, we nevertheless observed a modest but significant reduction in viral protein and RNA expression in infected 3T3 cells (Fig. 3). Finally, we asked whether overexpression of the key m^6^A reader YTHDF2, or downregulation of the m^5^C writer NSUN2, would affect MLV gene expression. Indeed, and as previously also reported for HIV-1 (8, 9), we observed enhanced MLV gene expression upon overexpression of murine YTHDF2 (Fig. 4A), and a substantial decline in MLV gene expression in 3T3 cells upon downregulation of NSUN2 expression using RNAi (Fig. 4C). These MLV data therefore confirm and extend our previously reported results, generated using HIV-1 (9) (8), indicating that m^6^A and m^5^C are positive regulators of viral gene expression and further argue that MLV, like HIV-1 and likely many other virus species, has evolved to use the epitranscriptomic writers and readers expressed by infected cells as a way to increase viral gene expression and replication.

It will therefore be of interest to investigate whether any viruses have also evolved the ability to upregulate the expression of these proteins.

## Materials & Methods

### Plasmids and cDNA cloning

A lentiviral vector was used to generate a clonal 3T3-derived cell line stably expressing FLAG-tagged mouse YTHDF2. Briefly, the mouse *ythdf2* gene (NP_663368) was PCR amplified from a cDNA library and cloned into the pLEX vector (9) 5’ to an internal ribosome entry site (IRES) and the puromycin (*puro*) resistance gene, all driven by the CMV immediate early promoter. A previously described (9) pLEX-based lentiviral vector expressing FLAG-tagged green fluorescent protein (GFP) was used as a control. A FLAG-tagged murine NSUN2 (NP_060225) expression plasmid was generated by PCR amplification of an NSUN2 cDNA that was then cloned into the pcDNA3.1 expression plasmid to generate pcDNA-FLAG-NSUN2. A mutant form of NSUN2 was generated by introducing the C271A mutation into pcDNA-FLAG-NSUN2. The NSUN2-C271A mutant spontaneously forms stable covalent bonds with target cytosines on RNA (6). The pcDNA-based expression plasmid ph3G-HA, expressing an HA-epitope tagged form of human APOBEC3G (A3G) has been described (29). Here, we substituted the FLAG epitope tag for HA to generate ph3G-FLAG.

The pNCS MLV proviral expression vector has been described and was a gift from Dr. Stephen Goff (18). Two MLV mutant clones were generated, one mutated to remove a single m^6^A site (Δm^6^A) and the second mutated to remove 4 m^5^C sites (Δm^5^C). Only silent mutations were introduced at these sites. Two DNA gBlocks were synthesized by IDT containing these silent mutations, and cloned into pNCS to generate pNCS-Δm^6^A and pNCS-Δm^5^C, respectively. Mutations introduced into pNCS-Δm^6^A are as follows; counting from the start codon of Env, with introduced mutations indicated in lower case letters: nt 354 5’-GAAGAgCCTctcACCTCC-3’.

Mutations introduced into pNCS-Δm^5^C are as follows; counting from the start codon of Gag, with introduced mutations again indicated in lower case letters: nt 2898 5’-GGtTTtTGTaGatTaTGGATt-3’, nt 3645 5’-agtGCTCAGaGaGCTGAAtTGATAGCAtTgACt-3’, nt 4242 5’-aGAACAtTaAAAAATATtACTGAGACtTGt-3’ and nt 4500 5’-ATtTTtCCtAGGTTt-3’. Clone integrity was verified by Sanger sequencing.

### MLV infections

To generate infectious MLV, pNCS-based proviral expression vectors (18) were transfected into 293T cells (CRL-3216; ATCC) using polyethyleneimine (PEI). At 24 h post-transfection, supernatant media were exchanged for fresh media. At 72 h post-transfection, the supernatant media containing infectious MLV virions were passed through a 0.45 μm filter and then used for infection of 3T3 cells.

### MLV gRNA purification

MLV virions were purified by a two-step method, as previously described (19). Briefly, the supernatant media from MLV-infected 3T3 cells were harvested at 72 hpi, passed through a 0.45 μm filter and the virions then pelleted through a 20% sucrose cushion by ultracentrifugation. The virion pellet was resuspended and layered onto a 7.2% to 20% iodixanol gradient (OptiPrep, Axis-Shield) prior to ultracentrifugation, to separate virions from cellular debris and exosomes. The virion band was then harvested and total RNA extracted using TRIzol. The isolated RNA was heat denatured in a loading buffer containing urea, and run on a preparative 1.5% TBE agarose gel. An RNA band of ~8kb, corresponding in size to the MLV gRNA, was visualized and excised and RNA isolated using acid phenol followed by phenol-chloroform extraction. The bulk of the purified MLV RNA was then used for UPLC-MS/MS analysis of RNA modifications while a small aliquot was retained for RNA-seq analysis, which was used to determine the purity of the MLV gRNA sample. RNA-seq was performed using the SMARTer^®^ Stranded Total RNA-Seq Kit v2 (NEB) following the manufacturer’s instructions.

### RNA modification identification by UPLC-MS/MS

Nucleosides were generated from purified MLV RNA by nuclease P1 digestion (Sigma) in buffer containing 25 mM NaCl and 2.5 mM ZnCl_2_ for 2 h at 37°C, followed by incubation with Antarctic Phosphatase (NEB) for an additional 2 h at 37°C (30). Nucleosides were separated and quantified using UPLC-MS/MS as previously described (17), except acetic acid was used in place of formic acid. Triplicate MLV gRNA samples were assessed by this method.

### PA-antibody-seq

PA-m^6^A-seq and PA-m^5^C-seq were performed as previously described (8–10). Briefly, 3T3 cells were infected with MLV as described above. At 48 hpi, cells were pulsed with 100 mM 4-thiouridine (4SU). After a further 24 h, total cellular RNA was extracted from the MLV-infected 3T3 cells using TRIzol, while MLV gRNA was extracted from virions that were collected by ultracentrifugation of the supernatant media through a 20% sucrose cushion. Total cellular poly(A)+ RNA was purified using oligo-dT magnetic beads (AM1922; Invitrogen) and 10 μg of poly(A)+ RNA or virion gRNA was then used following the previously reported PA-m^6^A-seq protocol (10, 24) using either an m^6^A-specific (202111; Synaptic Systems) or m^5^C-specific (C15200081; Diagenode) polyclonal antibody.

### NSUN2 packaging into virions

Plasmids expressing FLAG-tagged versions of the wild type or C271A mutant form of murine NSUN2, or human A3G, were co-transfected with pNCS into 293T cells using PEI. At 72 h post-transfection, the supernatant media were passed through 0.45μm filters and virions harvested by ultracentrifugation through a 20% sucrose cushion. Protein was extracted from the virion pellet in Laemmli buffer (31) before analysis by Western blot. Protein from producer cells was also harvested in Laemmli buffer to demonstrate protein expression from the transfected plasmids.

### Western blots

Proteins were extracted using Laemmli buffer, sonicated and denatured at 95°C for 10 min and then separated on Tris-Glycine-SDS polyacrylamide gels (Invitrogen). After electrophoresis, proteins were transferred to a nitrocellulose membrane, and then blocked in 5% milk in PBS + 0.1% Tween. Membranes were incubated in primary and secondary antibodies diluted in 5% milk in PBS + 0.1% Tween for 1 h each and then washed in PBS + 0.1% Tween. Each antibody was used at a 1:5000 dilution. The antibody targeting MLV Gag has been described (32) and was a gift from Dr. Stephen Goff. Antibodies targeting Actin (60008; Proteintech), NSUN2 (20854; Proteintech) and the FLAG epitope tag (F1804; Sigma), as well as anti-mouse HRP (A9044; Sigma) and anti-rabbit HRP (A6154; Sigma), were also used. Western blot signals were visualized by chemiluminescence. Image J was used for quantification of the intensity of protein bands.

### siRNA transfections

To investigate the effect of reduced NSUN2 protein levels on MLV protein expression, RNAi was utilized to knockdown NSUN2 expression in 293T cells. An siRNA specific to NSUN2 (siNSUN2), or a control siRNA (SR310319; Origene), was transfected into 293T cells at a concentration of 25 pmol/ml using Lipofectamine RNAiMAX (Invitrogen). At 48 h post-transfection, cells underwent a second siRNA transfection and were then incubated for a further 24 h. Cells were then transfected with pNCS. At 72 h post-transfection, the cells were harvested for Western blot analysis.

### YTDHF2 overexpression in 3T3 cells

The lentiviral expression vectors pLEX-GFP and pLEX-YTHDF2 were transfected into 293T cells along with the packaging plasmids pMD2G (12259; Addgene) and pΔ8.74 (22036; Addgene). Media were changed at 24 h post-transfection. At 72 h post-transfection the supernatant media containing the lentiviral particles was harvested and passed through a 0.45 μm filter and overlaid on 3T3 cells (CRL-1658; ATCC). At 48 hpi, transduced 3T3 cells were selected by the addition of puromycin to the culture media. After a further 72 h, cells were single cell cloned, expanded and assessed by Western blot for FLAG-GFP or FLAG-YTHDF2 expression. A single clone was then selected to determine the effect of overexpression of YTHDF2 on MLV infection. The GFP or YTHDF2 overexpressing cell lines were infected with MLV and at 72 hpi protein was harvested for Western blot analysis.

### Quantitative real-time PCR

Relative MLV gRNA expression levels were determined by qRT-PCR. The level of GAPDH mRNA was used to normalize all cellular qRT-PCR experiments, while 7SL RNA, which is packaged by MLV virions (20), was used to normalize virion qRT-PCR experiments. All primer sequences are listed below. RNA was extracted using the TRIzol method. cDNA was generated using the Ambion cDNA synthesis kit with random primers, following the manufacturer’s protocol. All qRT-PCR experiments were performed using Thermo Fisher’s Power Sybr Green PCR Master Mix (4367659; ABI) following the manufacturer’s instructions. All qRT-PCR data were quantified using the ΔΔCT method.

The primers sequences for GAPDH mRNA detection were as follows:

GAPDH Forward: 5’-TGGGTGTGAACCATGAGAAG-3’,

GAPDH Reverse: 5’-GATGGCATGGACT GTGGT C-3’,

The primers used for MLV genomic RNA detection were as follows:

MLV gag Forward: 5’-AGGAATAACACAAGGGCCCA-3’,

MLV gag Reverse: 5’-GGGTCCTCAGGGTCATAAGG-3’,

The primers used for human 7SL RNA detection were as follows:

7SL Forward: 5’GTGCGGACACCCGATCGGCA-3’,

7SL Reverse: 5’-TGAGGCTGGAGGATCGCTTGAG-3’.

### Bioinformatic analysis

Read alignments were performed using Bowtie (33). Reads were first aligned to the mouse genome, allowing up to 1 mismatch, then unaligned reads were aligned to the pNCS MLV transcriptome, again allowing up to 1 mismatch. At least one characteristic T to C mutation, resulting from 4SU incorporation and crosslinking to the antobody used, were required in both mouse and viral aligned reads. All data was processed using in-house Perl scripts and Samtools (34), and visualized with IGV, as previously described (10). The raw sequencing data obtained by RNA-seq have been submitted to the NCBI expression omnibus and are available under GenBank accession number GE (in process)

## Acknowledgments

This research was funded in part by NIH grants R01-DA046111, R56-AI124973 and U54-GM103297 to B.R.C. D.G.C. was funded by Marie-Skłodowska Curie Global Fellowship MSCA-IF-GF:747810. We thank the Duke Proteomics and Metabolomics Shared Resource, which performed mass spectrometry, and the Duke Center for Genomic and Computational Biology, which performed sequencing. We also thank Dr Stephen Goff for the gift of the pNCS MLV proviral expression plasmid and the MLV-Gag-specific antiserum.

D.G.C., A.C., H.P.B., B.A.L. and E.M.K. performed the experiments; D.G.C and E.M.K. analyzed the RNA-seq data; D.R.C. and B.R.C. wrote the manuscript and C.L.H. and B.R.C. oversaw the project.

## References

1. Meyer KD, Jaffrey SR. 2014. The dynamic epitranscriptome: N6-methyladenosine and gene expression control. Nat Rev Mol Cell Biol 15:313–326.

2. Roundtree IA, Evans ME, Pan T, He C. 2017. Dynamic RNA modifications in gene expression regulation. Cell 169:1187–1200.

3. Gilbert WV, Bell TA, Schaening C. 2016. Messenger RNA modifications: Form, distribution, and function. Science 352:1408–1412.

4. Yang X, Yang Y, Sun BF, Chen YS, Xu JW, Lai WY, Li A, Wang X, Bhattarai DP, Xiao W, Sun HY, Zhu Q, Ma HL, Adhikari S, Sun M, Hao YJ, Zhang B, Huang CM, Huang N, Jiang GB, Zhao YL, Wang HL, Sun YP, Yang YG. 2017. 5-methylcytosine promotes mRNA export - NSUN2 as the methyltransferase and ALYREF as an m5C reader. Cell Res 27:606–625.

5. Zhang X, Liu Z, Yi J, Tang H, Xing J, Yu M, Tong T, Shang Y, Gorospe M, Wang W. 2012. The tRNA methyltransferase NSun2 stabilizes p16INK(4) mRNA by methylating the 3’-untranslated region of p16. Nat Commun 3:712.

6. Hussain S, Sajini AA, Blanco S, Dietmann S, Lombard P, Sugimoto Y, Paramor M, Gleeson JG, Odom DT, Ule J, Frye M. 2013. NSun2-mediated cytosine-5 methylation of vault noncoding RNA determines its processing into regulatory small RNAs. Cell Rep 4:255–261.

7. Squires JE, Patel HR, Nousch M, Sibbritt T, Humphreys DT, Parker BJ, Suter CM, Preiss T. 2012. Widespread occurrence of 5-methylcytosine in human coding and non-coding RNA. Nucleic Acids Res 40:5023–5033.

8. Courtney DG, Tsai K, Bogerd HP, Kennedy EM, Law BA, Emery A, Swanstrom R, Holley CL, Cullen BR. 2019. Epitranscriptomic Regulation of HIV-1 Gene Expression by m5C and the Novel m5C Reader MBD2. Available at SSRN: https://ssrn.com/abstract=3334977.

9. Kennedy EM, Bogerd HP, Kornepati AV, Kang D, Ghoshal D, Marshall JB, Poling BC, Tsai K, Gokhale NS, Horner SM, Cullen BR. 2016. Posttranscriptional m(6)A editing of HIV-1 mRNAs enhances viral gene expression. Cell Host Microbe 19:675–685.

10. Courtney DG, Kennedy EM, Dumm RE, Bogerd HP, Tsai K, Heaton NS, Cullen BR. 2017. Epitranscriptomic enhancement of influenza A virus gene expression and replication. Cell Host Microbe 22:377–386 e375.

11. Tsai K, Courtney DG, Cullen BR. 2018. Addition of m6A to SV40 late mRNAs enhances viral structural gene expression and replication. PLoS Pathog 14:e1006919.

12. Lichinchi G, Gao S, Saletore Y, Gonzalez GM, Bansal V, Wang Y, Mason CE, Rana TM. 2016. Dynamics of the human and viral m6A RNA methylomes during HIV-1 infection of T cells. Nature Microbiology 1:16011.

13. Hao H, Hao S, Chen H, Chen Z, Zhang Y, Wang J, Wang H, Zhang B, Qiu J, Deng F, Guan W. 2019. N6-methyladenosine modification and METTL3 modulate enterovirus 71 replication. Nucleic Acids Res 47:362–374.

14. Ye F, Chen ER, Nilsen TW. 2017. Kaposi’s Sarcoma-Associated Herpesvirus Utilizes and Manipulates RNA N6-Adenosine Methylation To Promote Lytic Replication. J Virol 91.

15. Hesser CR, Karijolich J, Dominissini D, He C, Glaunsinger BA. 2018. N6-methyladenosine modification and the YTHDF2 reader protein play cell type specific roles in lytic viral gene expression during Kaposi’s sarcoma-associated herpesvirus infection. PLoS Pathog 14:e1006995.

16. Ringeard M, Marchand V, Decroly E, Motorin Y, Bennasser Y. 2019. FTSJ3 is an RNA 2’-O-methyltransferase recruited by HIV to avoid innate immune sensing. Nature 565:500–504.

17. Basanta-Sanchez M, Temple S, Ansari SA, D’Amico A, Agris PF. 2016. Attomole quantification and global profile of RNA modifications: Epitranscriptome of human neural stem cells. Nucleic Acids Res 44:e26.

18. Gao G, Goff SP. 1998. Replication defect of moloney murine leukemia virus with a mutant reverse transcriptase that can incorporate ribonucleotides and deoxyribonucleotides. J Virol 72:5905–5911.

19. Bogerd HP, Kennedy EM, Whisnant AW, Cullen BR. 2017. Induced packaging of cellular microRNAs into HIV-1 virions can inhibit infectivity. mBio 8:e02125–02116.

20. Onafuwa-Nuga AA, King SR, Telesnitsky A. 2005. Nonrandom packaging of host RNAs in moloney murine leukemia virus. J Virol 79:13528–13537.

21. Li Q, Li X, Tang H, Jiang B, Dou Y, Gorospe M, Wang W. 2017. NSUN2-Mediated m5C Methylation and METTL3/METTL14-Mediated m6A Methylation Cooperatively Enhance p21 Translation. J Cell Biochem 118:2587–2598.

22. Li X, Xiong X, Yi C. 2016. Epitranscriptome sequencing technologies: decoding RNA modifications. Nat Methods 14:23–31.

23. Hafner M, Landthaler M, Burger L, Khorshid M, Hausser J, Berninger P, Rothballer A, Ascano M, Jr., Jungkamp AC, Munschauer M, Ulrich A, Wardle GS, Dewell S, Zavolan M, Tuschl T. 2010. Transcriptome-wide identification of RNA-binding protein and microRNA target sites by PAR-CLIP. Cell 141:129–141.

24. Chen K, Lu Z, Wang X, Fu Y, Luo GZ, Liu N, Han D, Dominissini D, Dai Q, Pan T, He C. 2015. High-resolution N(6)-methyladenosine (m(6) A) map using photo-crosslinking-assisted m(6) A sequencing. Angew Chem Int Ed Engl 54:1587–1590.

25. Liu B, Merriman DK, Choi SH, Schumacher MA, Plangger R, Kreutz C, Horner SM, Meyer KD, Al-Hashimi HM. 2018. A potentially abundant junctional RNA motif stabilized by m(6)A and Mg(2). Nat Commun 9:2761.

26. Tirumuru N, Zhao BS, Lu W, Lu Z, He C, Wu L. 2016. N(6)-methyladenosine of HIV-1 RNA regulates viral infection and HIV-1 Gag protein expression. Elife 5.

27. Gokhale NS, McIntyre AB, McFadden MJ, Roder AE, Kennedy EM, Gandara JA, Hopcraft SE, Quicke KM, Vazquez C, Willer J, Ilkayeva OR, Law BA, Holley CL, Garcia-Blanco MA, Evans MJ, Suthar MS, Bradrick SS, Mason CE, Horner SM. 2016. N6-methyladenosine in Flaviviridae viral RNA genomes regulates infection. Cell Host Microbe 20:654–665.

28. Lichinchi G, Zhao BS, Wu Y, Lu Z, Qin Y, He C, Rana TM. 2016. Dynamics of human and viral RNA methylation during Zika virus infection. Cell Host Microbe 20:666–673.

29. Bogerd HP, Doehle BP, Wiegand HL, Cullen BR. 2004. A single amino acid difference in the host APOBEC3G protein controls the primate species specificity of HIV type 1 virion infectivity factor. Proc Natl Acad Sci U S A 101:3770–3774.

30. Dominissini D, Nachtergaele S, Moshitch-Moshkovitz S, Peer E, Kol N, Ben-Haim MS, Dai Q, Di Segni A, Salmon-Divon M, Clark WC, Zheng G, Pan T, Solomon O, Eyal E, Hershkovitz V, Han D, Dore LC, Amariglio N, Rechavi G, He C. 2016. The dynamic N(1)-methyladenosine methylome in eukaryotic messenger RNA. Nature 530:441–446.

31. Laemmli UK. 1970. Cleavage of structural proteins during the assembly of the head of bacteriophage T4. Nature 227:680–685.

32. Rodriguez JJ, Goff SP. 2010. Xenotropic murine leukemia virus-related virus establishes an efficient spreading infection and exhibits enhanced transcriptional activity in prostate carcinoma cells. J Virol 84:2556–2562.

33. Langmead B, Trapnell C, Pop M, Salzberg SL. 2009. Ultrafast and memory-efficient alignment of short DNA sequences to the human genome. Genome Biol 10:R25.

34. Li H, Handsaker B, Wysoker A, Fennell T, Ruan J, Homer N, Marth G, Abecasis G, Durbin R, Genome Project Data Processing S. 2009. The Sequence Alignment/Map format and SAMtools. Bioinformatics 25:2078–2079.

